# High throughput RNA sequencing discovers symptomatic and latent viruses: an example from ornamental Hibiscus

**DOI:** 10.1101/2022.01.25.477650

**Authors:** Shuyu Zhou, Katja R. Richert-Pöggeler, Ziwei Wang, Trude Schwarzacher, J.S. (Pat) Heslop-Harrison, Qing Liu

## Abstract

*Hibiscus rosa-sinensis* L. (Hibiscus, Malvaceae) is an ornamental species grown widely in amenity plantings. We collected leaves on an urban roadside pavement (sidewalk) near a market in Guangzhou which showed multiple symptoms of leaf rolling, deformation and chlorosis. Initial evaluation by electron microscopy using negative staining of drip preparations revealed the presence of tobamovirus-like particles. Total RNA was extracted, and, unusually, without any RNA selection based on sequence, was used for cDNA library construction and high-throughput survey sequencing. From the 814 Mb of clean sequence data (from 2,712,161 paired reads of 150 bp) reads representing chloroplast, ribosomal, and mitochondrial genes were filtered out, eliminating 79.1% of reads. 1,135,848 × 150 bp of the sequence was retained and screened for viral sequences. Assembly of these sequences detected nine virus species from seven virus genera comprising three tobamoviruses, namely, Tobacco mosaic virus, Tobacco mild green mosaic virus and Hibiscus latent Singapore virus, Turnip mosaic virus (*Potyvirus*), Potato virus M (*Carlavirus*), Hibiscus chlorotic ringspot virus (*Betacarmovirus*), Fabavirus sp (*Fabavirus*), Cotton leaf curl Multan virus (*Begomovirus*) and a putative mitoviruses replicating in mitochondria, *Chenopodium quinoa* mitovirus 1. Mapping the reads to complete virus reference sequences showed high and uniform coverage of the genomes from 3,729 x coverage for Turnip mosaic virus to 22 x for Cotton leaf curl Multan virus. By comparison, nuclear reference genes actin showed 14 x coverage and polyubiquitin 27 x. Notable variants from reference sequences (SNPs) were identified. With the low cost of sequencing and potential for semi-automated bioinformatic pipelines, the whole-RNA approach has huge potential for identifying multiple undiagnosed viruses in ornamental plants, resulting in the ability to take preventive measures in production facilities against spread and to product quality for the mutual benefit of producers and consumers.

## INTRODUCTION

*Hibiscus rosa-sinensis* (*H. rosa-sinensis*)is a common ornamental plant in subtropical and tropical areas, including China, where the species is widely planted as single specimens, in hedges, or in landscaping, in gardens and for amenity horticulture, due to their dense growth and attractive flowers (Adkins et al., 2006). Hibiscus extracts are also used for tea, and in skin ointments with traditional healing uses (Ryoo et al., 2010; Tsai, 2016). Related species are used as textile fibres (kenaf) and planted in temperate gardens. Both RNA and DNA viruses infect *H. rosa-sinensis* worldwide (Sastry et al., 2019) including alfamo-betacarmo, rhabdo-, tobamo and begomoviruses (Table 1) where they may reduce quality and cause economic losses.

**Table 1.** Taxonomy, genome type, virion morphology, distribution and year of 1^st^ report of selected viruses infecting *Hibiscus rosa-sinensis*. Those found in the RNA sequencing study here are in bold.

| Species | Genus, Family | Genome | Morphology | Distribution | Transmission | 1<sup>st</sup>Report | Reference |
| --- | --- | --- | --- | --- | --- | --- | --- |
| Alfalfa mosaic virus | Alfamovirus, Bromoviridae | +ssRNA | bacilliform (spherical) | Spain | Aphids, sap | 2012 | Parrella et al. |
| Citrus leprosis virus | Cilevirus, Kitaviridae | +ssRNA | bacilliform (spherical) | Hawaii | Mites, sap | 2013 | Melzer et al. |
| Cotton leaf curl Kokhran virus | Begomovirus, Geminiviridae | ssDNA | isometric | Pakistan | whitefly | 2014 | Akhtar et al. |
| **Cotton leaf curl Multan virus** | **Begomovirus, Geminiviridae** | **ssDNA** | **isometric** | **China, Philippines, India** | **whitefly** | **2005** | **Rajeshwari et al.** |
| Eggplant mottled dwarf nucleorhabdovirus | Nucleorhabdovirus, Rhabdoviridae | -ssRNA | bacilliform | Canary islands, Yugoslavia, Morocco, Spain, Italy, Greece | Leafhopper | 2008 | De Stradis et al. |
| **Hibiscus chlorotic ringspot virus** | **Betacarmovirus, Tombusviridae** | **+ssRNA** | **isometric** | **Iran, Australia, El Salvador, Fidji, Italy, the Solomon Islands, Thailand, Taiwan, New Zealand, Nigeria, Singapore, Malaysia, Bangladesh and USA** | **Mechanical inoculation** | **1980** | **Jones and Behncken** |
| Hibiscus latent<br>Fort Pierce virus | Tobamovirus,<br>Virgaviridae | +ssRNA | rod-shaped | USA, Italy and Japan | Mechanical<br>inoculation, sap | 2003 | Adkins et<br>al. |
| Hibiscus latent<br>ringspot virus | Nepovirus,<br>Secoviridae | +ssRNA | isometric | Australia, USA, Italy, Bangladesh,<br>Nigeria | Mechanical<br>inoculation, sap | 1980 | Brunt et al. |
| **Hibiscus latent**<br>**Singapore**<br>**virus** | **Tobamovirus,**<br>**Virgaviridae** | **+ssRNA** | **rod-shaped** | **Singapore, Taiwan and Italy** | **Mechanical**<br>**inoculation,**<br>**sap** | **2005** | **Srinivasan**<br>**et al.** |
| Impatiens<br>necrotic spot<br>orthotospovirus | Orthotospovirus,<br>Tospoviridae | +/-ssRNA | enveloped,<br>spherical | New Zealand | Thrips, sap | 2009 | Elliott et al. |
| Okra mosaic<br>virus | Tymovirus,<br>Tymoviridae | +RNA | isometric | Ivory coast, Nigeria |  | 1972 | Givord et<br>al. |
| Tomato mosaic<br>virus | Tobamovirus,<br>Virgaviridae | +ssRNA | rod-shaped | China | Mechanical<br>inoculation, sap | 2004 | Huang et al. |

Virus-infected *H. rosa-sinensis* can display various symptoms. Infection with Hibiscus chlorotic ringspot virus (HCRSV) exhibits mottling or chlorotic spots, rings, or vein-banding symptoms under natural conditions, which may disappear during the summer months (Jones and Behncken, 1980). Infection with Cotton leaf curl Multan virus (CLCuMuV), exhibits leaf curling, vein swelling, vein darkening and enations on the veins on the undersides of the leaves (She et al., 2017). Visual in/spection of symptomatic phenotypes needs verification by additional analysis since symptom expression can be influenced by various abiotic and biotic parameters. Symptoms may also vary (and typical symptoms may be modified) in plants with mixed infections, and expression can vary depending on the cultivar.

Plant virology and virus diagnostics use multiple technologies including bioassays and serological testing (ELISA), electron microscopy and PCR methods (RT-PCR, qPCR) as well as transmission experiments for virus characterization. In 2009, high-throughput sequencing technology (HTS) entered the field of plant virology (Kreuze et al., 2009) and it has been a valuable addition to plant virus detection and identification tools, along with the reference sequences in databases such as GenBank. Like electron microscopy, HTS does not need previous knowledge on the viruses’ infection characteristics, nor specific antibodies or sequence amplification primers. The International Committee on Taxonomy of Viruses (ICTV) has recognized the use of metagenomic sequencing data and applied it to virus taxonomy (Simmonds et al., 2017). As an example, infection of Hibiscus latent Fort Pierce virus in *H. rosa-sinensis* was first identified by the combination of routine diagnostic tools and RNA-seq method in China (Lan et al., 2020), and conventional methods, like RT-PCR, were used to verify the sequencing results, demonstrating the reliability of HTS for analyzing the virus genome sequence.

In this study, we aimed to detect viruses by high-throughput survey sequencing of RNA, without previous knowledge of the viruses present or their infection characteristics. Analyzing the viral genome sequence and establishing a rapid and sensitive virus detection method is of great significance for the prevention and control of viral diseases.

## MATERIALS AND METHODS

### Plant material

Symptomatic leaves of *H. rosa-sinensis* were sampled from plants growing in soil squares on an urban roadside pavement (US: sidewalk) near Changban market, Guangzhou, China (23º10’E, 113º21’N, Xingke Rd/Lu, Tianhe Qu, 23.174231N, 113.361287). Fresh leaves from a single plant, collected without contamination, were rinsed with water and used for RNA extraction.

### Transmission electron microscopy

A mixed sample of symptomatic leaves was homogenized in 2–5-fold volume of extraction buffer (0.1 M phosphate buffer pH 7.0 containing 2 % polyvinylpyrrolidone MW 11.000 and 0.2 % sodium sulfite). Viral particles were adsorbed by floating a pioloform carbon-coated copper grid on the leaf extract. For negative contrast the grids were stained with 1 % uranyl acetate in ultrapure water, dried and analysed by transmission electron microscopy (TEM) in a Tecnai G2 Spirit TWIN Transmission Electron Microscope (Czech Republic) with 80 kV accelerating voltage. Digital images were acquired with a GATAN camera and processed using TIA software (Tecnai Imaging and Analysis).

### Extraction of the total RNA and sequencing library construction

Total RNA was extracted from leaves with Monarch Total RNA Miniprep Kit (#T2010S, New England Biolabs), following the manufacturer’s instructions. The extracted RNA was used for library preparation (NEBNext Ultra II RNA Library Prep with Sample Purification Beads, #E7775S) with appropriate barcoded sequencing primers (NEB #E7335-12 indices).

### High-throughput sequencing, assembly, and analysis of viral sequences

The cDNA library was sequenced by Wuhan Bena Technology Service using Illumina Hiseq X-ten (2×150 bp paired-end with 350 bp insert size average). A total of 814 Mb of sequence with the appropriate barcode primers was extracted.

The paired-end reads were assembled to known abundant reference sequences of the Hibiscus (https://www.ncbi.nlm.nih.gov/nuccore/?term=) before *de novo* assembly of the remaining reads (Geneious assembler, Medium-low sensitivity/Fast setting). Contig sequences were analysed using BLAST comparisons (National Centre for Biotechnology Information (NCBI) database). Where the sequence showed homology to known viral genomes, these were used as a reference target for reassembly of the reads as used for the preliminary assembly (Figure 1).

**Figure 1.**
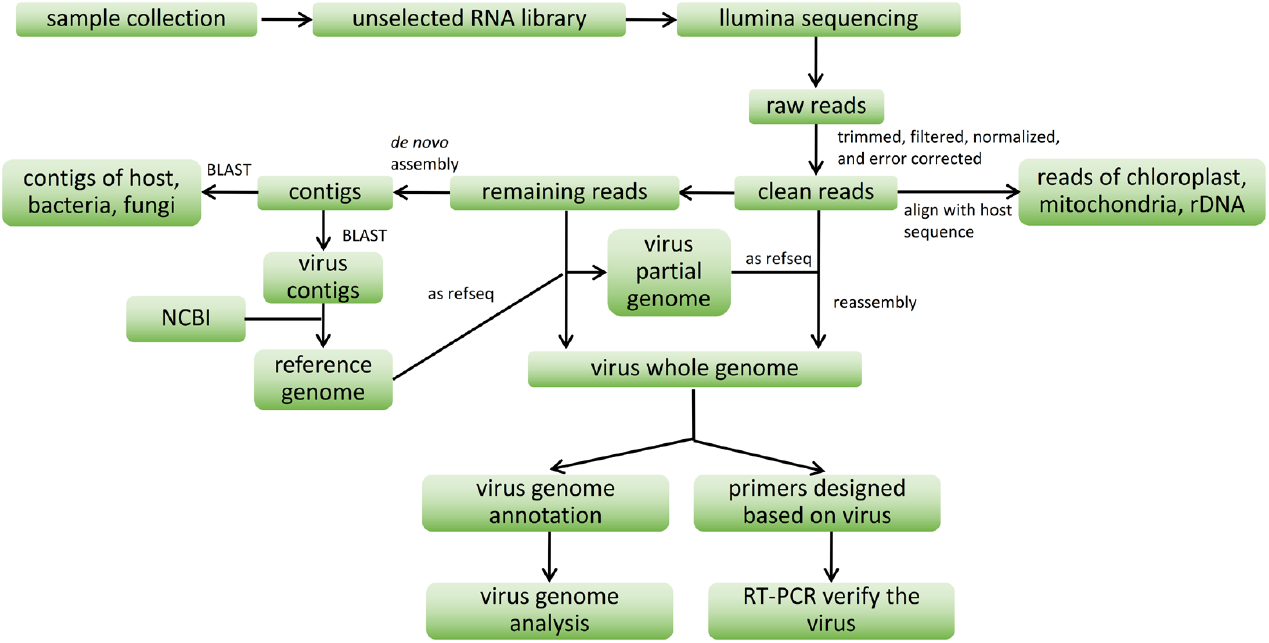
The flow-chart used for analysis of raw reads from RNA and bioinformatics.

### Sequence comparisons and phylogenetic analyses

The virus complete genome was aligned with those of reference sequences published in NCBI, using Clustal W sequence alignment. Phylogenetic trees were generated by the MAGE X (Kumar, 2018) software with 1,000 bootstrap replicates.

## RESULTS AND DISCUSSION

### Identification of viruses in *Hibiscus rosa-sinensis* samples

Leaves of *H. rosa-sinensis* showing leaf curl and chlorosis (Figure 2A) indicated probable virus infection. Symptomatic leaves were analysed for the presence of infectious entities using electron microscopy (Richert-Pöggeler et al., 2019). Negative staining of drip preparations revealed multiple rod-shaped virions resembling those of tobamoviruses in length and width (Figure 2B & Figure 2C). Thus, these results confirmed that *H. rosa-sinensis* is infected by virus.

**Figure 2.**
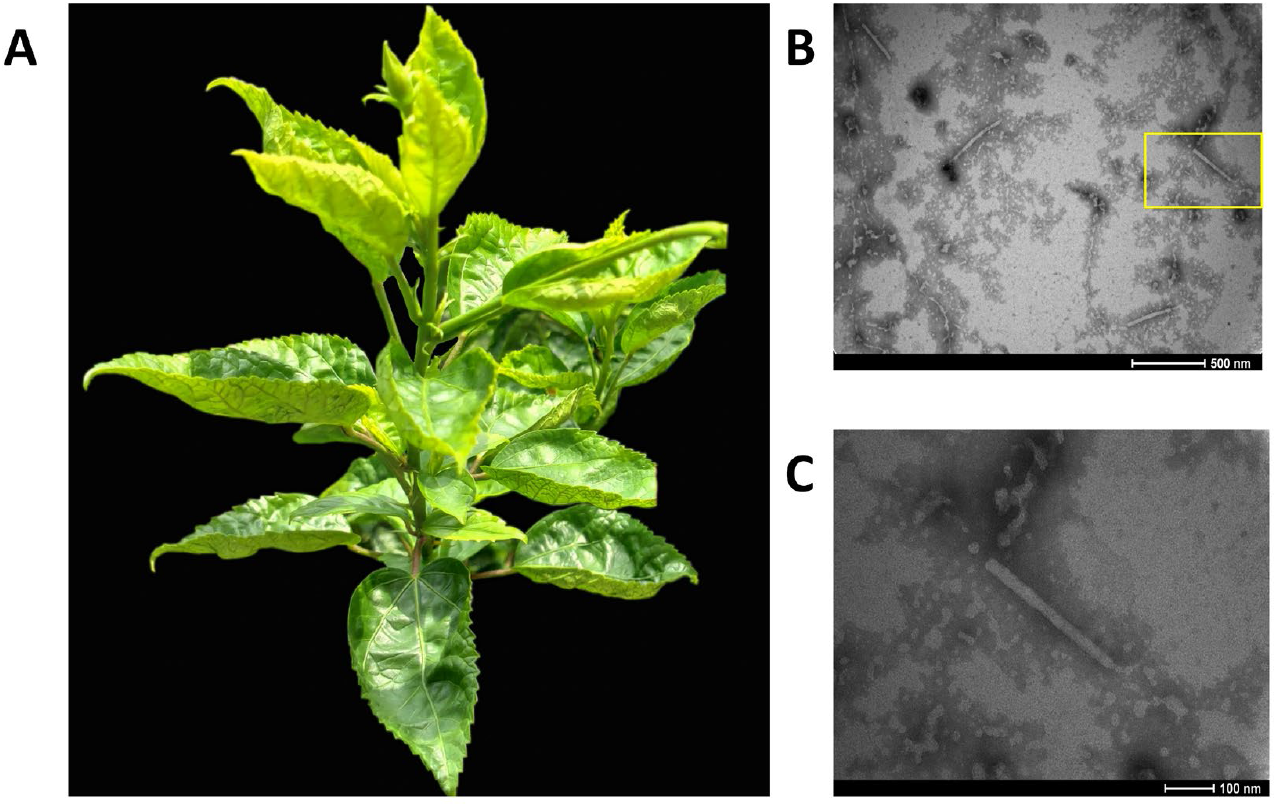
Detection of virus infection in *Hibiscus rosa-sinensis* sampled from a roadside planting near Changban Market, Guangzhou, China. (A) Symptoms of leaves infected with viruses. (B) Electron micrographs of homogenized and negative stained leaves from (A) showing tobamovirus-like virions. Scale bar = 500 nm. (C) Magnification of a single virion circled by yellow box from (B) with rod-shaped morphology of about 300 nm in length and 18 nm in width, scale bar = 100 nm.

### Sequencing, assembly, and identification of viral sequences

The unselected RNA library from *H. rosa-sinensis* had a total of 5,424,322 high-quality reads. After mapping to the rDNA sequence (KX709342), chloroplast genome (MK937807) and mitochondrial genome (NC_035549), 20.9% of clean reads remained. The proportion of rDNA, chloroplast and mitochondria removed were 51.0%, 26.1% and 1.1% respectively, plus 0.9% of related unpaired reads. *De novo* assembly of the remaining reads (“screened RNA reads”), generating 19,776 contigs, including Hibiscus nuclear transcripts as well as viral RNA sequences. Contigs were compared to the NCBI database by BLAST and viral genomes were identified. The screened DNA reads were then assembled to these reference viral sequences.

A total of nine abundant virus genomes were identified in the samples: Cotton leaf curl Multan virus (CLCuMuV), *Chenopodium quinoa* mitovirus 1 (CqMV1), Fabavirus sp., Hibiscus chlorotic ringspot virus (HCRSV), Hibiscus latent Singapore virus (HLSV), Potato virus M (PVM), Turnip mosaic virus (TuMV), Tobacco mosaic virus (TMV) and Tobacco mild green mosaic virus (TMGMV). The number of reads representing each virus differed widely, with coverage of the genome ranging from 22 x (CLCuMuV, 0.008% of the reads) to 3,596 x (TuMV, 4.35% of the reads) (Figure 3). The screened RNA sequences include nuclear gene transcripts, and the presence and abundance of two constitutively expressed genes often used as standards for gene expression (RNA transcription) were analysed. Of the 5,424,322 reads, 71 were assembled (14 x coverage) to *Hibiscus cannabinus* actin (ACT) mRNA (DQ866836) and 192 were assembled (27 x coverage) to *Hibiscus syriacus* polyubiquitin mRNA (XM_039204349).

**Figure 3.**
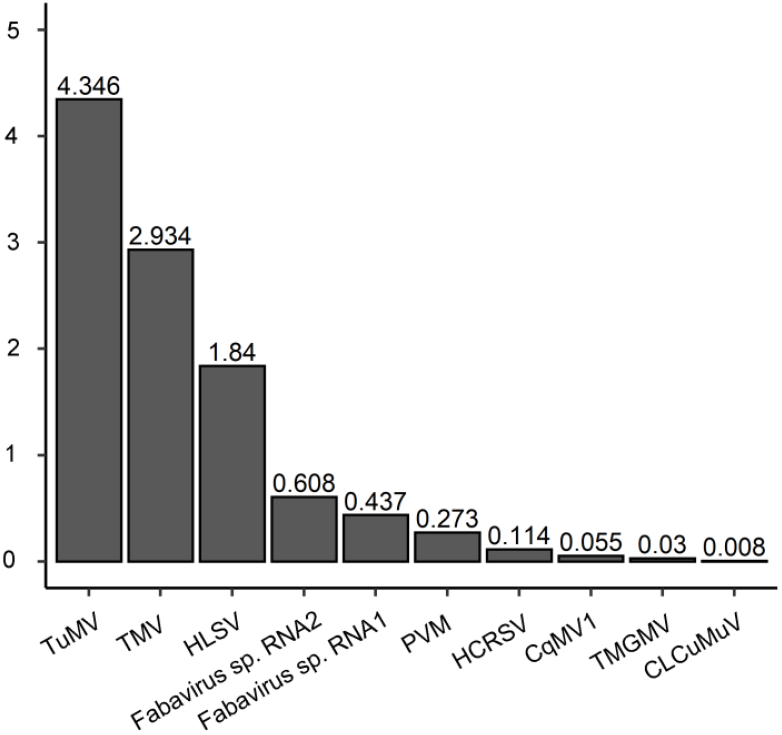
The proportion of screened RNA reads mapping to genomes of nine viruses of *Hibiscus rosa-sinensis* detected in the RNA reads (Virus abbreviations as Table 2.)

Mapping the reads to reference genomes showed relatively even coverage of the complete genome sequences (Figure 4). Notably, one domain of the CLCuMuV isolate (from bp470 to 620 representing part of the coat protein AV1, encoded from bp276 to 1046) was not present in *H. rosa-sinensis* sequencing RNA data. 5’ and 3’ ends of each virus were not accurately defined, and ends sometimes showed degenerate or shared domains; further work is required to define both ends of the new viral sequences.

**Table 2.** Genbank reference sequences with highest identity with virus sequences isolated from *Hibiscus rosa-sinensis* RNA.

| **Virus isolates** | **Abbreviation** | **Identity** | **Accession number** | **Year** | **Original place** | **Reference** |
| --- | --- | --- | --- | --- | --- | --- |
| Chenopodium quinoa mitovirus 1 | CqMV1 | 99.8% | NC_040543 | 2016 | Italy | Direct Submission |
| Cotton leaf curl Multan virus | CLCuMuV | 97.9% | JQ424826 | 2013 | China | Tang et al., 2013 |
| Fabavirus sp. RNA1 | Fabavirus | 98.1% | MN253486 | 2017 | China | Direct Submission |
| Fabavirus sp. RNA2 | Fabavirus | 96.6% | MN253487 | 2017 | China | Direct Submission |
| Hibiscus chlorotic ringspot virus | HCRSV | 95.0% | KC876666 | 2012 | Israel | Luria et al., 2013 |
| Hibiscus latent Singapore virus | HLSV | 91.5% | MZ020960 | 2017 | China | Direct Submission |
| Potato virus M | PVM | 79.6% | MT114149 | 2018 | Canada | Direct Submission |
| Tobacco mild green mosaic virus | TMGMV | 98.5% | KM596785 | 2013 | China | Direct Submission |
| Tobacco mosaic virus | TMV | 98.0% | HE818416 | 2012 | China | Yang et al., 2012 |
| Turnip mosaic virus | TuMV | 98.5% | LC537509 | 2014 | China | Kawakubo et al., 2021 |

**Figure 4.**
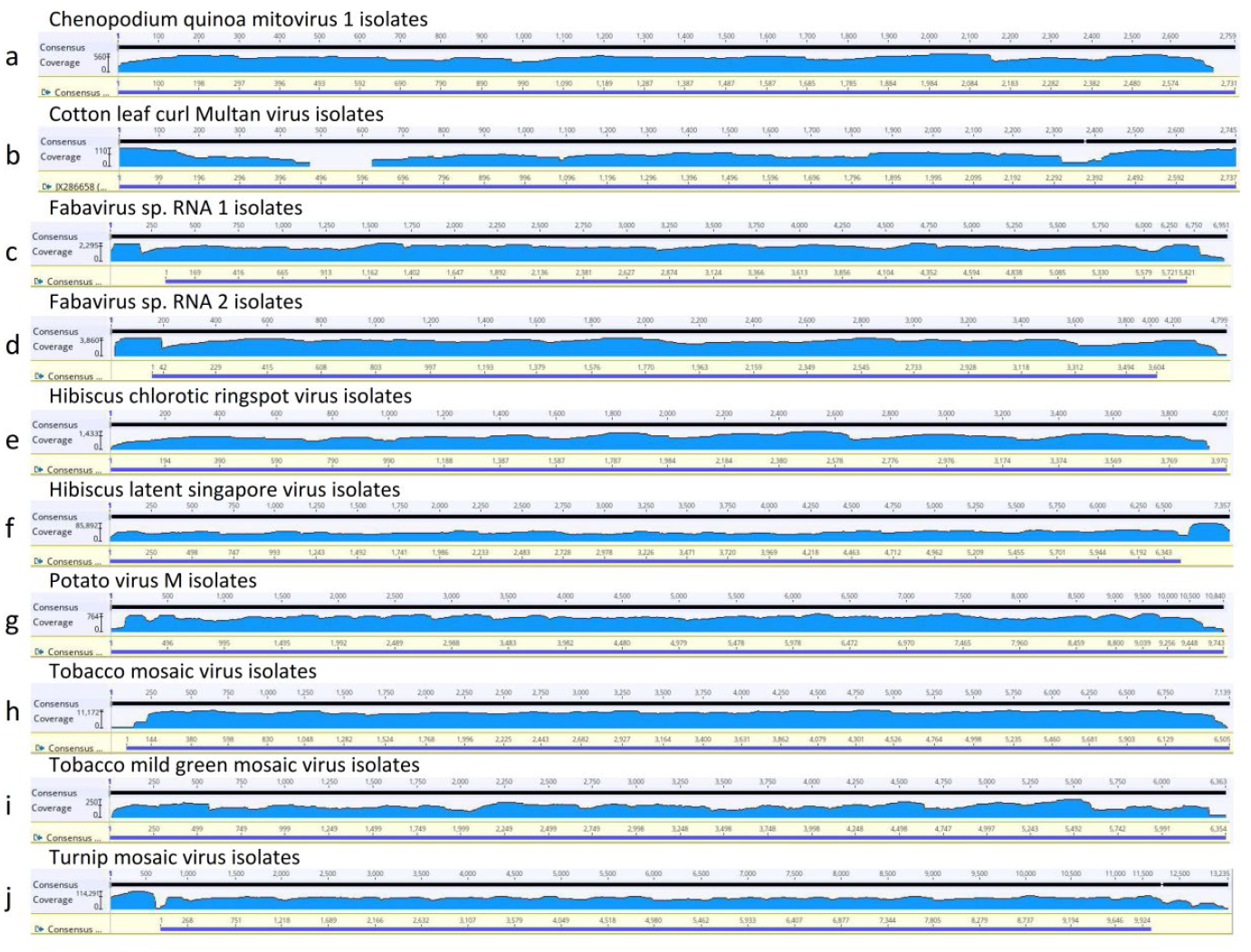
Assembly of high-throughput unselected RNA reads from *H. rosa-sinensis* to complete reference virus sequences identified in the RNA data. (a) CqMV1 isolate Che1 (NC_040543); (b) CLCuMuV isolate GD11 (JQ424826); (c) Fabavirus sp. RNA1 isolate BJ (MN253486); (d) Fabavirus sp. RNA2 isolate BJ (MN253487); (e) HCRSV isolate SBO1 (MK279671); (f) HLSV isolate HLSV-UKM (MN080499); (g) PVM isolate K (MT114149); (h) TMV isolate Fumeng (HE818416); (i) TMGMV isolates HN (KM596785); (j) TuMV isolate CHN313 (LC537509).

A phylogenetic analysis (Figure 5) was carried out for the well-characterised TMGMV using 16 related TMGMV isolates (Table 3) sequences deposited in GenBank. The sequence identified here groups most closely with a TMGMV isolate from *Capsicum annuum* (chilli) found in the Hunan province, in central and southern China (Table 3).

**Table 3.** Accession number, origin and host plant of Tobacco mild green mosaic virus (TMGMV) isolates used in the phylogenetic analysis (figure 5).

| Accession number | Serial number | Place | Host | Time | Reference |
| --- | --- | --- | --- | --- | --- |
| AB078435 | J | Japanese | Nicotiana benthamiana | 2002 | Morishima et al. |
| DQ821941 | HP | Taiwan | Capsicum annuum | 2006 | Direct Submission |
| EF469769 | DSMZ PV-0113 | USA | Columnea allenii x Columnea crassifolia | 2007 | Direct Submission |
| HRO1_TMGMV | HRO1_TMGMV | China | Hibiscus | 2019 | This study |
| JX534224 | Xiamen | China | Capsicum annuum | 2013 | Direct Submission |
| KM596785 | HN | China | Capsicum annuum | 2014 | Direct Submission |
| MF139550 | TN29 | China | Capsicum annuum | 2017 | Direct Submission |
| MH730970 | Ng92/73 | Spain | Nicotiana glauca | 2018 | Bera et al. |
| MK005155 | TMGMV-CaJO | Jordan | Pepper | 2019 | Favara et al. |
| MK648158 | WW17-E1-Nocc-neg | Slovenia | Nicotiana occidentalis | 2019 | Font et al. |
| MN267902 | WW17-E1-Nocc-pos | Slovenia | Nicotiana occidentalis | 2019 | Direct Submission |
| MN267903 | DSMZ PV-0112 | USA | Nicotiana tabacum | 2020 | Direct Submission |
| MT675965 | NBHP | Vietnam | Pepper | 2020 | Direct Submission |
| MW012408 | DSMZ PV-0124 | Italy | Nicotiana tabacum | 2021 | Choi et al. |
| MZ405634 | TMGMVgp1 | USA | Nicotiana tabacum | 1990 | Direct Submission |
| NC_001556 | BR | Brazil | Nicotiana glauca | 2018 | Solis et al. |

**Figure 5.**
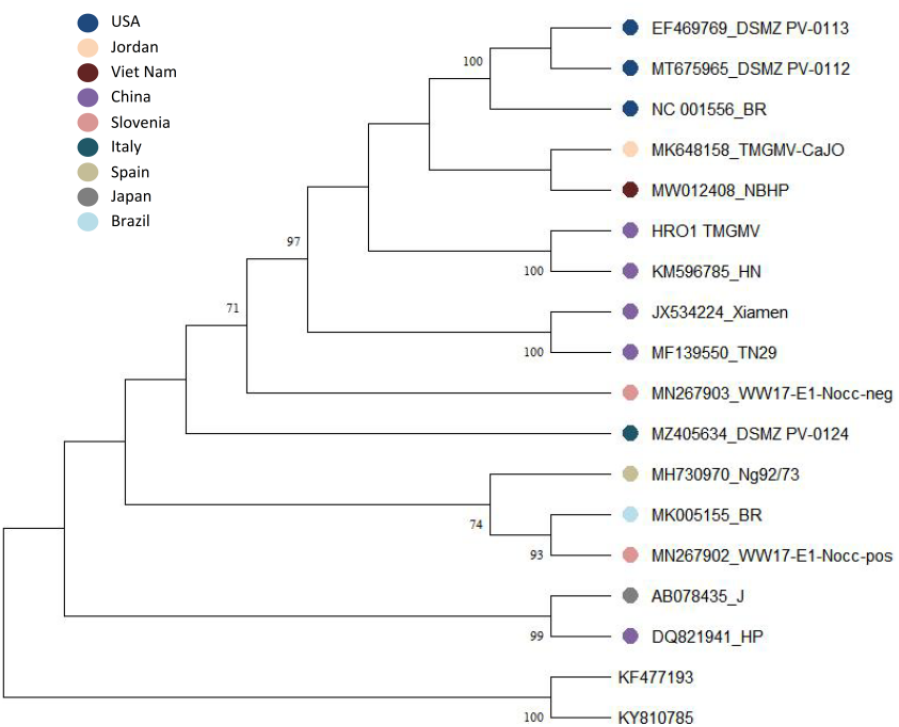
Tobacco mild green mosaic virus (TMGMV) neighbor-joining phylogenetic tree based on whole genome sequences of related TMGMV isolates deposited at GenBank and out of this study, Tomato mottle mosaic virus (KF477193) and Tobacco mosaic virus (KY810785) being outgroups, with 1,000 bootstrap replicates.

## CONCLUSIONS

The unselected, total RNA libraries proved suitable for detection of viral genomes including +ssRNA and ssDNA viruses (Table 2, Figure 3), as well as single (tobamo-, poty-, mito-, betacarmovirus) and multicomponent (alfamo-, begomo-, fabavirus) genomes. The filtering method (Figure 1) successfully separated host from virus sequences, and the huge volume of RNA sequence obtained meant that discarding 79.1% of sequences, representing the major RNA species (chloroplast, mitochondria and rRNA), and including the nuclear genome transcripts in the analysis, did not compromise analysis of the viral sequences. The comprehensive approach captured sequences from all cellular compartments such as nucleus, mitochondria and cytoplasm, including viruses replicating in cytoplasm and nucleus/cytoplasmic compartments. We identified mixed infection in *H. rosa-sinensis* with nine viruses, including a potyvirus (TuMV) and a tobamovirus (TMV) as highly abundant copies – up to 138 times the expression of the actin nuclear reference gene.

Combining results from EM and HTS we can conclude that tobamoviruses are extremely abundant. Immuno-electron microscopy (IEM) will be a valuable tool to investigate if virions are produced from all viruses identified in HTS. We note the higher amount of fabavirus RNA2 sequences coding for the capsid protein compared to fabavirus RNA1 encoding for the replicase. RT-PCR approaches would also be useful to test the presence of the identified viruses in additional plant accessions. Both quantitative RT-PCR and small RNA sequencing will be useful tools to further study the interferences of identified viruses in the mixed infection with each other and the host respectively. Nevertheless, compared to the unselective library construction and moderate cost of high-throughput sequencing (c. $USD200 per sample), both the high researcher effort and time required for establishing RT-PCR protocols for detection of multiple viruses, and the costs of sequencing multiple individual PCR products (10 PCR products per virus, 9 viruses, so 100 sequencing reactions at c. $USD5), mean that RT-PCR will be at least as expensive as whole RNA sequencing used here. Arguably, the high-throughput results are more accurate with fewer experimental steps and no primer selection.

The data reveal two cases for expansion of the host range of viruses (Table 1 vs Table 2): for TuMV from Brassicales (turnip) to Malvales (*H. rosa-sinensis*), that are phylogenetically related (APG IV, 2016), and for the fabavirus from Asterales (*Dahlia* sp.) to Malvales that are more distantly related. In several virus sequences, the most similar references were from Chinese virus isolates from other species (families; eg *Capsicum*, Solanaceae), suggesting recent and most likely ongoing transfer between plant families. We did not detect potyvirus-like virions with helical structure and filamentous morphology using electron microscopy because virion concentration in the sample most likely was below the detection limit.

The biological impact of all identified viruses should be confirmed in bioassay testing, followed by IEM where antisera are available. The 1000-fold range in virus abundance (Figure 3) would be interesting to put into context with participation in both infection and symptom expression. The consequences of the virus presence will be especially important for risk assessment for ornamental plants, where viruses such as PVM are known to be latent (Richert-Pöggeler et al., 2015; Chofong et al., 2021), and planting material may be transported over long distances.

Finally, the identification of Chenopodium quinoa mitovirus 1 contributes to the knowledge about abundance of mitoviruses and their impact on horizontal DNA transfer (Hillmann et al., 2018).

Like many ornamental species selected by horticulturalists, Hibiscus thrives and flowers in public plantings under the high virus load revealed here, where there are also many abiotic stresses (including water and proximity to people or traffic). Knowledge of virus presence may enable more vigorous plants with less infection or higher resistance to be selected, and reduce the opportunities for virus transfer to natural environments. High-throughput RNA sequencing is an important approach for non-selective and efficient enumeration of the viral load in ornamental species.

## ACKNOWLEDGEMENTS

The authors want to thank Rufang Deng for taking the transmission electron microscopy images of the virus. This work was supported by grants from National Science Foundation of China (32070359), Guangdong Basic and Applied Basic Research Foundation (2021A1515012410), Overseas Distinguished Scholar Project of SCBG (Y861041001) and Undergraduate Innovation Training Program of Chinese Academy of Sciences (KCJH-80107-2020-004-97). KRP received a travel grant from the German Ministry of Agriculture in frame of the Bilateral Scientific Cooperation and Exchange Project between Germany and China (323-06.01-03-11/2018-CHN).

